# Assembly-free single-molecule nanopore sequencing recovers complete virus genomes from natural microbial communities

**DOI:** 10.1101/619684

**Authors:** John Beaulaurier, Elaine Luo, John Eppley, Paul Den Uyl, Xiaoguang Dai, Daniel J Turner, Matthew Pendelton, Sissel Juul, Eoghan Harrington, Edward F. DeLong

## Abstract

Viruses are the most abundant biological entities on Earth, and play key roles in host ecology, evolution, and horizontal gene transfer. Despite recent progress in viral metagenomics, the inherent genetic complexity of virus populations still poses technical difficulties for recovering complete virus genomes from natural assemblages. To address these challenges, we developed an assembly-free, single-molecule nanopore sequencing approach enabling direct recovery of high-quality viral genome sequences from environmental samples. Our method yielded over a thousand high quality, full-length draft virus genome sequences that could not be fully recovered using short read assembly approaches applied to the same samples. Additionally, novel DNA sequences were discovered whose repeat structures, gene contents and concatemer lengths suggested that they represent phage-inducible chromosomal islands that were packaged as concatemers within phage particles. Our new approach provided novel insight into genome structures, population biology, and ecology of naturally occurring viruses and viral parasites.

## Introduction

Viruses are central to the ecology and evolution of virtually all cellular lifeforms on Earth. Viruses impact microbial diversity and activity by exerting top-down control on dominant microorganisms, releasing organic matter from cell lysis, and accelerating genetic exchange via horizontal gene transfer. Due to their importance in early molecular genetic studies and their comparatively small genome size, viruses were the first biological entities whose genomes were fully sequenced (Fiers et al. 1976; Sanger et al. 1977). More recently, application of microbial community shotgun genome sequencing (metagenomics) has advanced understanding of virus populations in the wild (Breitbart et al. 2002; Hurwitz et al. 2013; Sullivan 2015). Metagenomic studies to date have uncovered thousands of novel viral genes, genomic fragments, and genomes from the oceans (Mizuno et al. 2013a; Brum et al. 2015; Roux et al. 2016b; Aylward et al. 2017; Luo et al. 2017; Roux et al. 2018).

The ecological richness, evenness and genomic complexity of virus assemblages still complicates determination of full-length virus genome sequences from naturally occurring viral populations. Previous viral metagenomic studies have relied primarily on three main approaches: 1. Whole metagenomic short-read shotgun sequencing and assembly (Breitbart et al. 2002; Hurwitz et al. 2013; Sullivan 2015) 2. Fosmid-based large DNA insert shotgun sequencing (Mizuno et al. 2013a); 3. Amplification-based shotgun sequencing approaches, including single cell multiple-displacement amplification single-cell techniques, and PCR-based linker ligation amplification (Hurwitz et al. 2013; Roux et al. 2014). Each approach, despite its individual advantages, also has its own unique limitations, which can result from amplification biases, challenges associated with De Bruijn graph short read assemblies, and the limited DNA insert size range and cloning biases associated with fosmids. Due to these difficulties, obtaining whole virus genome sequences from complex naturally occurring populations remains a challenge.

Given the typical size range of double-stranded DNA viruses infecting bacteria and archaea (∼3-300 kb), we reasoned that determining entire viral genome sequences from single reads should possible using single-molecule nanopore sequencing approaches. Here we describe and assess a method for obtaining whole virus genome sequences from naturally occurring populations using Oxford Nanopore Technologies’ (ONT) platform. The ONT platform enables the sequencing of a single molecule of DNA by measuring changes in electrical current as it passes through a nanopore, and, from those measurements, inferring deoxynucleotide base sequence composition. The method was tested and validated on virus-enriched aquatic samples. We show here that a single sequencing run in an amplification-free library prepared from virus-enriched high molecular weight DNA can generate over 1000 high-quality polished draft genomes, that we refer to here as assembly-free virus genomes (AFVGs), from complex environmental samples. We show these AFVGs can provide new insight into the diversity and nature of viruses and viral parasites in natural microbial populations.

## Results

### Virus particle collection and DNA yields

Viral particles collected from 82 L of 15 m seawater by pre-filtration though 0.2 μm filters, and final recovery on 0.03 μm filters, yielded a total of 1.7 μg of purified high molecular weight DNA. Viral particles collected from 100 L of 0.1 μm pre-filtered seawater that was subsequently concentrated via tangential flow filtration, yielded a total of 5.3 μg and 3.5 μg of purified high molecular weight DNA from the 117 m and 250 m seawater samples, respectively.

### ONT Sequencing

A total of 1 μg of purified DNA from each sample was used to prepare sequencing libraries using ONT’s standard ligation sequencing kit LSK109, modifying the DNA repair and end-prep incubation step to 20 min. Sequencing was conducted on GridION X5 with FLO-MIN106 (R 9.4.1) flowcells (Oxford Nanopore). The sequencing output from each of the three flowcells ranged from 5.15 - 12.28 gigabases, generating read lengths up to 254 kb (Table 1). The 15 m sample produced the most reads, although these were generally shorter (12.9 kb read N50) than those obtained from the 117 m and 250 m samples (29.0 and 38.0 kb read N50, respectively).

**Table 1.** Sequencing, binning, and polished genome statistics for virus-enriched seawater samples obtained from three different depths. During the binning procedure, a subset of the 5-mer bins were enriched for reads of a particular length. Each of these bins consists of one or more average nucleotide identity (ANI) clusters. Draft genomes were generated by polishing a single representative read from each ANI cluster and were subsequently deduplicated across the “cellular” and “non-cellular” read partitions to ensure that only unique polished genomes were obtained from each depth.

| Sample |  | Sequencing statistics |  |  | Binning statistics |  |  | Genome statistics |  |
| --- | --- | --- | --- | --- | --- | --- | --- | --- | --- |
| Depth | Annotation | # reads | Bases (Gb) | Read N50 (Kb) | 5-mer bins | Read length enriched bins | ANI clusters | # unique polished genomes | Mean genome length (Kb) |
| 15 m | Cellular | 536,625 | 3.88 | 14.32 | 137 | 23 | 14 | 24 | 41.33 |
|  | Non-cellular | 465,096 | 1.82 | 9.09 | 109 | 48 | 45 |  |  |
| 117 m | Cellular | 113,661 | 2.11 | 30.81 | 319 | 121 | 133 | 176 | 48.87 |
|  | Non-cellular | 227,687 | 3.04 | 27.6 | 475 | 203 | 221 |  |  |
| 250 m | Cellular | 267,036 | 7.12 | 38.1 | 1074 | 940 | 1330 | 1430 | 41.64 |
|  | Non-cellular | 291,241 | 5.16 | 37.96 | 890 | 827 | 1216 |  |  |
| uvMED | Simulated | 386,000 | 2.89 | 18.7 | 279 | 191 | 187 | 178 | 36.64 |

### Identification of marine virus genomes from metagenomic nanopore reads

An assembly-free bioinformatic pipeline was developed to isolate and polish full-length phage genomes from nanopore reads (Fig. 1). Reads were first binned into two partitions using Kaiju (Menzel et al. 2016) (Table 1) to separate cellular reads (prokaryotic and eukaryotic) from remaining reads (viral and unclassified). Subsequent read binning and genome polishing for each depth sample was conducted separately within each read partition. Briefly, the pipeline consisted of representing each read by a vector of normalized 5-mer frequencies, using dimensionality reduction and clustering tools to embed these 5-mer frequency vectors in two dimensions and call read bins, clustering reads within each bin based on pairwise average nucleotide identity (ANI), and polishing a single read from each ANI cluster with the other reads in the cluster (see Methods).

**Figure 1.**
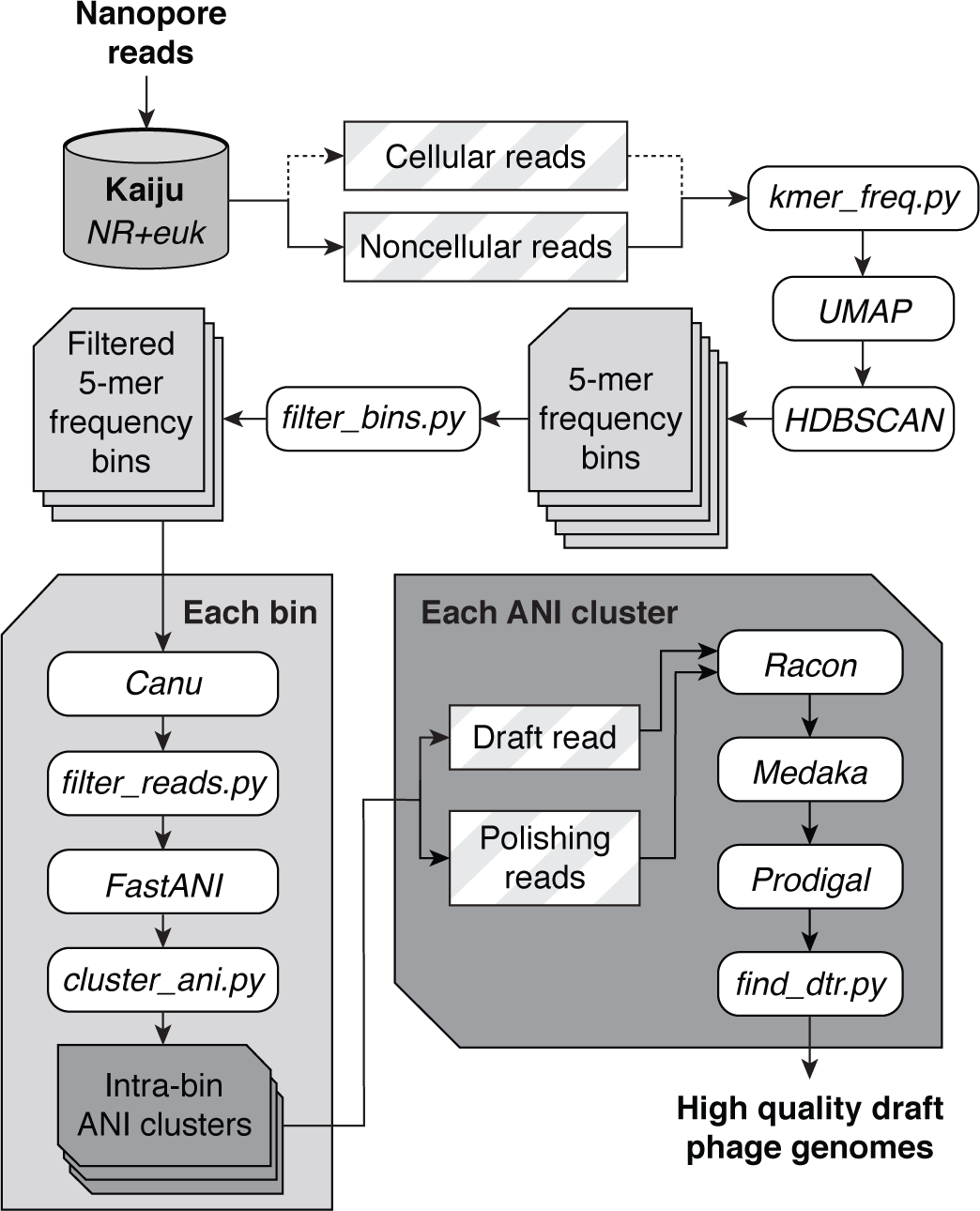
Bioinformatic pipeline for assembly-free discovery of marine phage genomes in nanopore sequences. Nanopore reads from each depth sample were first separated into “cellular” and “noncellular” partitions based on their Kaiju annotations (Menzel et al. 2016). Read sequences were then decomposed into vectors containing their normalized 5-mer frequencies, then all read vectors were embedded in a 2-dimensional map using the dimensionality reduction tool UMAP (McInnes et al. 2018). Bins were called from the embedding using HDBSCAN (McInnes et al. 2017) and filtered to identify bins enriched for genome-scale read lengths. For each remaining bin, reads were error corrected using Canu (Koren et al. 2017) and genome-scale reads were selected. FastANI (Jain et al. 2018) was used to calculate pairwise average nucleotide identity (ANI) values for all genome-scale reads. Finally, a representative read was selected from each ANI cluster and polished by the remaining reads in the ANI cluster using a combination of consensus polishing tools. Each polished draft genome was subsequently annotated for protein coding sequences using Prodigal (Hyatt et al. 2010).

We first validated the phage discovery pipeline on a set of 193 known marine phage genomes from the uvMED collection (Mizuno et al. 2016). A total of 2,000 simulated nanopore reads with realistic length and error profiles were generated for each genome, then combined to form a mock viral metagenome. Our bioinformatic pipeline resulted in 279 5-mer frequency bins (Supplementary Fig. S1), of which 191 were found to contain bin-specific peaks in their read length distribution indicating the presence of full-length viral reads (Table 1). These bins resulted in 187 total ANI bin clusters, 178 of which produced a unique polished draft genome. Comparison of the polished draft genomes with the original reference sequences revealed that 174/193 original genomes were recapitulated at > 99% identity and >99% genome coverage. Recapitulation of the remaining genomes is made difficult by the fact that several uvMED genomes appear to be circular permutations of each other, making it difficult to differentiate them using global sequence approximations like 5-mer frequency and ANI.

We next applied the phage discovery pipeline to the three virus-enriched seawater samples recovered from depths of 15 m, 117 m, and 250 m. The assembly-free nature of the phage genome discovery pipeline allowed us to isolate and polish genomes containing complex repeat structures that can be problematic for short read metagenomic assemblies. For example, an 11.9 kb complex repeat structure in genome AFVG_250N00740 was easily resolved by selecting one 67.3 kb phage read from the 5-mer bin ANI cluster and polishing it with the remaining reads from the same cluster, each of which fully spanned the complex repeat (Supplementary Fig. S2). Furthermore, direct terminal repeats (DTR), consisting of identical sequences of up to 16 kb that flank linear virus genomes, are characteristic of many dsDNA tailed phages (Casjens and Gilcrease 2009). We used the DTRs to confirm that reads indeed spanned a full virus genome, and for initial virus characterization reported here, all polished sequences lacking a DTR were discarded. We observed DTRs in the polished draft genomes between 31 bp to 6.1 kb in length. Notably, such repeats would not be detected in short read assembly approaches, as they would either collapse into a single copy or produce circularly permuted mis-assemblies.

Additionally, the phage genome discovery pipeline preserved levels of microdiversity that are known to induce fragmentation in short-read viral metagenomic assemblies (Roux et al. 2017). For example, genomes AFVG_250N00862 and AFVG_250N01202 were obtained from reads found in a single 5-mer bin (Fig. 2a). This bin was highly enriched for reads of ∼35 kb (Fig. 2b), suggesting that they either derived from the same phage genome or from closely related genomes. Hierarchical clustering of pairwise ANI calculations separated the reads into two distinct ANI clusters, suggesting the existence of two populations of phage that differ at the strain level (Fig. 2c). Subsequent comparison between the two polished genomes obtained from these ANI clusters revealed sequences of >96 % ANI totaling more than 17 kb, 9 kb of which shares >99% ANI, interspersed with multi-kb regions of significant sequence divergence (Fig. 2d).

**Figure 2.**
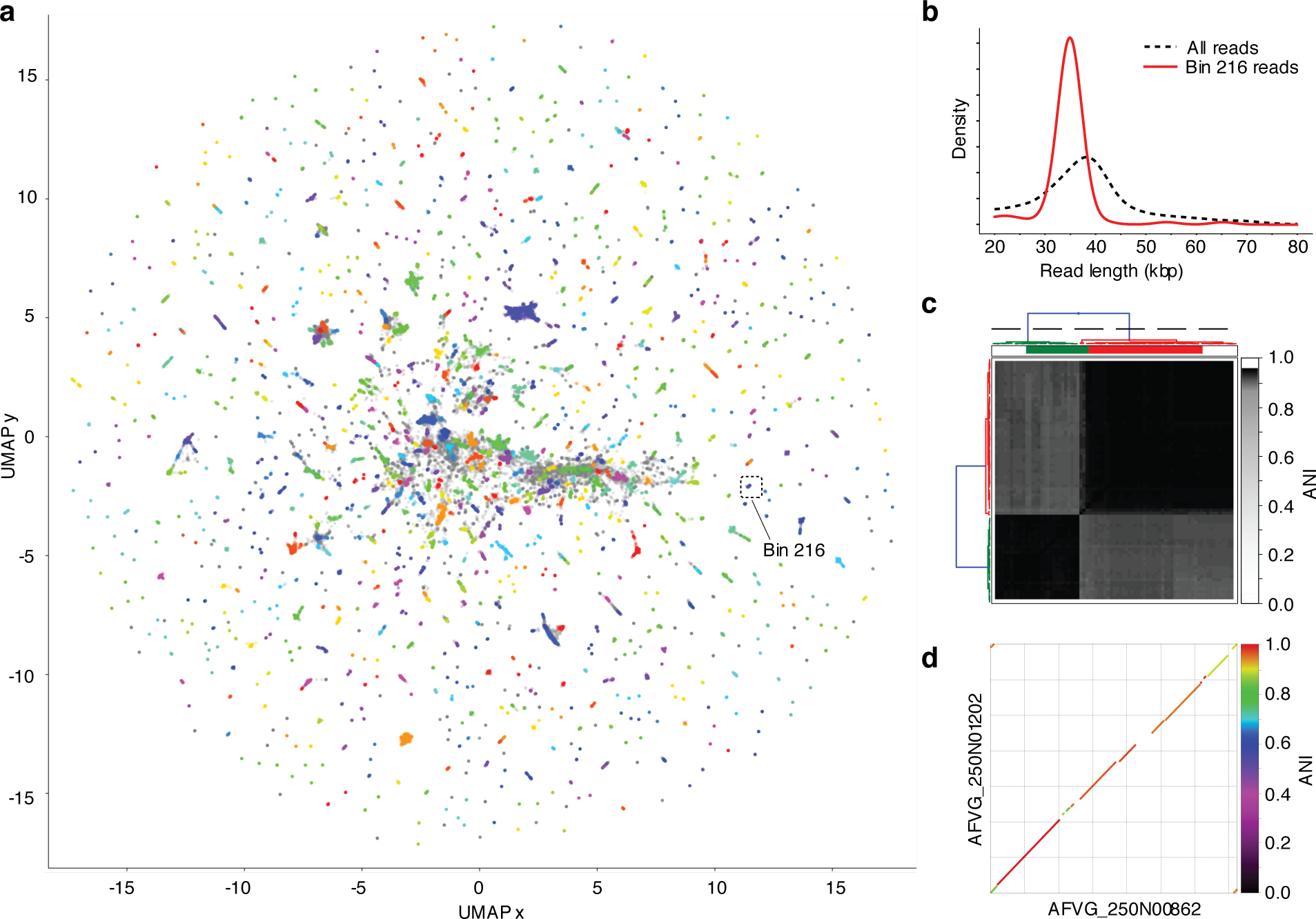
Phage discovery steps for 250 m sample. (**a**) Nanopore reads longer than 15 kb and annotated bioinformatically as “noncellular” were first represented by their normalized 5-mer frequencies and dimensionally reduced into a 2D embedding using UMAP (McInnes et al. 2018). Reads in the 2D embedding are colored based on their assignment to the 890 bins called by HDBSCAN (McInnes et al. 2017). Some bin colors are redundant due to the large number of bins. Reads not assigned to a bin are colored grey. (**b**) Read lengths from bin 216 were compared to read lengths from all bins, revealing a notable enrichment of ∼35 kb reads and a depletion of reads at other lengths. (**c**) Among these genome-scale reads in bin 216, hierarchical clustering of pairwise average nucleotide identity (ANI) values revealed two ANI clusters from which a single polished phage genome was generated for each (AFVG_250N01202 and AFVG_250N00862). (**d**) Comparison between the AFVG_250N01202 and AFVG_250N00862 shows microdiversity between the two polished phage genomes. Large regions of high nucleotide sequence identity (several at >95%) are interspersed with regions of divergent sequence.

The number of unique AFVGs obtained from each sample was elevated in samples from greater water depths (Table 1). The 15 m sample (which used the filter collection method and a 0.2 μm prefilter) produced 24 polished draft genomes, while the 117 m and 250 m samples yielded 176 and 1430 unique AFVGs, respectively (Supplementary Fig. S3). The higher yield of very long reads in the 250 m sample compared with the 15 m sample undoubtedly affected this result, although the larger number of observed 5-mer frequency bins at lower depths also suggested an increase in phage sequence diversity.

The yields of different viral genotypes from short-read Illumina sequencing versus nanopore sequencing in libraries prepared from the same DNA was compared for all three samples (Supplementary Fig. S4). In each of the three samples, the coverage of Illumina short reads of any given AFVG was comparable to the number of nanopore reads within that same AFVGs k-mer bin. While there were a very few outliers with higher representation in the short-read libraries (Supplementary Fig. S4), in general the recovery of sequences scaled similarly in the short read versus long-read nanopore sequence datasets prepared from the same samples. The nanopore sequencing approach described here recovered many more complete virus genome sequences than did short read sequencing and assembly approaches alone, using DNA from the same samples. More specifically, all the AFVGs produced by our nanopore sequencing method were longer (and more complete) compared to any homologous contigs recovered in the short-read sequence assemblies from the identical DNA sample (Supplementary Fig. S5a). Conversely, any short read contigs having homology to the nanopore AFVGs from the same DNA sample, were all fully covered by the nanopore AFVGs (Supplementary Fig. S5b).

### Validating AFVG origins and preliminary characterization

The assembly-free virus genomes (AFVGs) produced by our method were evaluated and validated using a variety of well-established approaches for characterizing viral metagenomic sequences (Roux et al. 2015; Bolduc et al. 2017; Hurwitz et al. 2018; Roux et al. 2018). The primary k-mer binning (Fig. 2a) generated nanopore sequence clusters having discrete and tight sequence length distributions, that were very different from bulk read size distributions (Fig. 2b; Supplementary Table S1). We postulated that these bin-specific tight read length distributions, ranging between 28.0 – 87.3 kb, indicated that we recovered assembly-free full-length viral genomes contained in single nanopore sequence reads. The three samples from different depths varied with respect to AFVG size ranges (Supplementary Fig. S3; Supplementary Table S1). Specifically, AFVGs in the 15 m sample ranged from 31.1 – 69.7 kb (average 41.3 kb), in the 117 m sample from 32.9 – 87.3 kb (average 48.9 kb), and in the 250 m sample from 28.0 – 72.1 kb (average 41.6). These values are comparable to previously reported planktonic viral isolate and community genome size distributions (Steward et al. 2000; Holmfeldt et al. 2013). No AFVGs larger than about 90 kbp were detected in our current assembly-free nanopore sequencing method, since there were few sequence reads >100 kbp.

To further validate the viral origins of the AFVGs, we examined the putative virus genome sequences identified in our bioinformatic pipeline for potential virus signatures (Roux et al. 2015; Hurwitz et al. 2018), specific viral gene content, and similarity to global viral gene sequence databases (Hurwitz and Sullivan 2013; Mizuno et al. 2013b; Brum et al. 2015; Paez-Espino et al. 2016; Roux et al. 2016a). All of the 1630 AFVGs identified in our nanopore sequence pipeline were also flagged as viral by the virus sequence classifier Virsorter (Roux et al. 2015). Since a final selection criterion in our pipeline included requiring DTRs be present in the putative AFVGs (a characteristic of many linear double stranded bacteriophage genomes (Casjens and Gilcrease 2009)), all or our AFVGs contained this viral feature. The DTRs (ranging between 31 - 6108 bp in length) had average lengths of 227 bp, 995 bp, and 518 bp in the 15 m, 117 m and 250 m samples, respectively (Supplementary Table S1).

Although few of the protein coding sequences in the AFVGs were bore high similarity to those in NCBI’s RefSeq database (Supplementary Fig. S6), a high proportion of AFVG annotated genes did share high sequence similarities to virus genes found in global metagenomic viral databases (Fig. 3; (Roux et al. 2015; Bolduc et al. 2017; Hurwitz et al. 2018; Roux et al. 2018). The majority of individual AFVGs had >60% of their annotated genes matching at >60% average amino acid identity (AAI) to protein coding virus genes in viral reference databases. AFVGs in the 250 m sample contained the largest proportion of genes that did not match any of those in available databases (Fig. 3), in congruence with previous findings (Luo et al. 2017). The AFVGs also contained a high proportion of viral marker genes, including genes annotated as terminase, tail, head, capsid, portal and integrase proteins (Supplementary Fig. S7). These virus marker genes also tended to decrease in their overall proportions in AFVGs from the 250 m sample (Supplementary Fig. S7), again reflecting the less well characterized nature of deep-sea viral populations.

**Figure 3.**
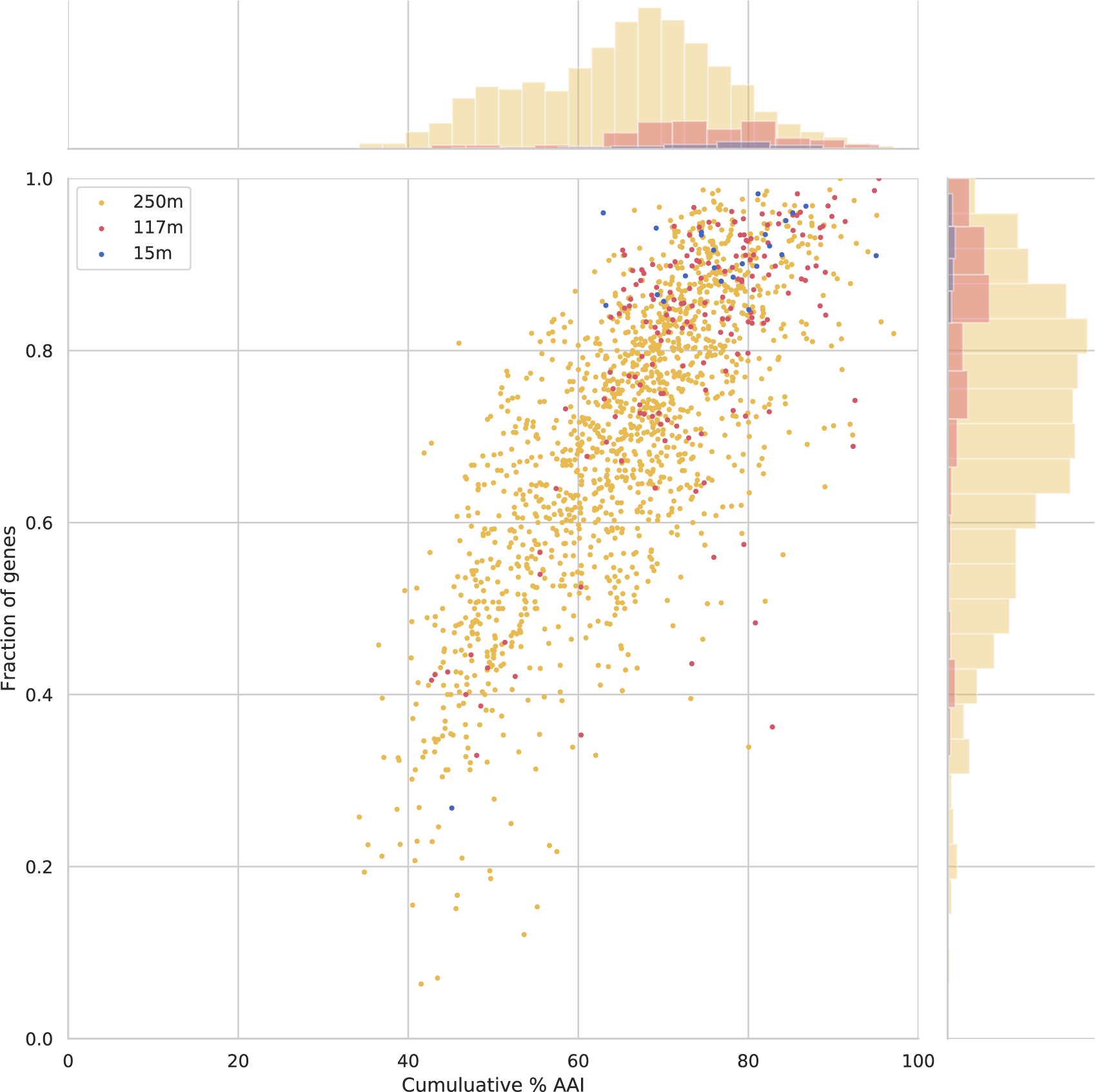
Similarity of AFVG annotated genes to known viral genes. Each point represents an AFVG colored by the depth at which it was found. The y-axis encodes the fraction of genes with matches to known viral genes (y-axis), and the x-axis encodes the cumulative percent amino acid identity (%AAI) of those matches. The marginal histograms show the distribution of values for “cumulative %AAI” (top) and “fraction of genes” (right) grouped by depth.

The putative taxonomic affiliations of AFVGs were consistent with the known microbial community composition and depth distributions of their planktonic microbial hosts from the same oceanographic region from which our samples originated (Supplementary Fig. S8; (Delong et al. 2006; Pham et al. 2008; Aylward et al. 2017; Luo et al. 2017; Mende et al. 2017)). In particular a large proportion of AFVGs bore genes that were highly similar to those from cyanophage, *Pelagibacter* (SAR11) phage, *Puniceispirillum* (SAR116) phage, as well as viruses resembling those that infect *Pseudomonas* and *Vibrio* species (Supplementary Fig. S9 a,b,c). Consistent with individual marker gene results, AFVGs from the deepest sample (250 m) shared the least similarity to known and well-characterized virus types.

### Linear concatemer sequences isolated from seawater

Among the polished AFVGs, eleven were 34.3-38.1 kb sequences composed exclusively of concatemers of 5.4-7.8 kb repeat sequences (Supplementary Table S2). To further explore this unexpected phenomenon, we searched for linear concatemers among all reads from each of the three samples. In the 117 m sample, 854 linear >15 kb concatemer reads had a significant enrichment of read lengths between 60-65 kb (Fig. 4a). We determined the repeat copy number in the concatemeric reads by dividing the read length by the length of the repeated sequence unit, and found that many of the reads contained whole integer numbers of repeats (between 5 and 7) with no partial repeat copies on either end (Fig. 4b). Interestingly, 41/176 (23%) of the AFVGs produced from the 117 m sample were larger than 60 kb, as opposed to only 77/1430 (5%) for the 250 m sample (Supplementary Fig. S3). In the 250 m sample, 1879 linear >15 kb concatemer reads were found with a significant read-length enrichment between 35-40 kb (Fig. 4c). These concatemer size distributions were generally consistent with the (non-concatemeric) AFVGs obtained from 250 m, where 1148/1430 (80%) of the AFVGs were shorter than 45 kb (Supplementary Fig. S3). In concordance with the 117 m sample, we found that many concatemeric reads from the 250 m sample contained a whole integer number of repeats (between 4-7 whole repeat copies) with no partial repeat copies on either end (Fig. 4d).

**Figure 4.**
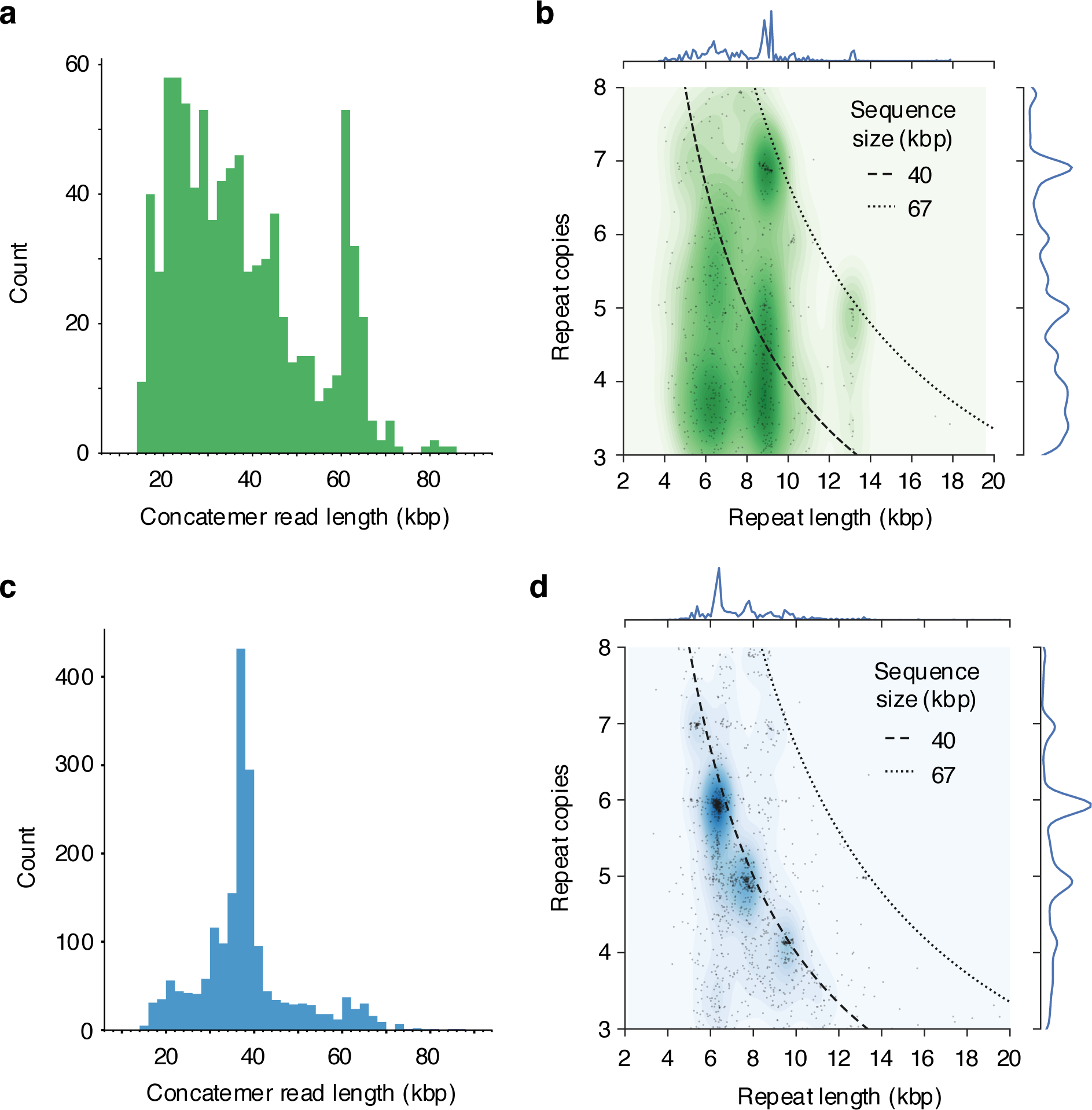
Read lengths and repeat copy numbers in concatemeric nanopore reads. The length distribution of linear concatemer reads isolated from the 117 m sample shows an anomalous spike in concatemer read lengths between 60-65 kb. These concatemer read lengths are similar in size to many of the complete phage genomes obtained from the 117 m sample. (**b**) The length of the repeat and the number of repeats in the concatemer sequence showing an enrichment of concatemer reads having a whole number of repeat copies (most notably at 5 and 7 copies) in the 117 m sample. (**c**) Concatemeric read length distribution from the 250 m sample showing a strong enrichment for concatemer reads between 35-40 kb. (**d**) The pattern of concatemer reads containing whole number copy repeats is even more pronounced in the 250 m sample, where the majority of reads are shorter than 40 kb and contain between 4-7 whole repeat copies.

Annotation of the eleven polished concatemer sequences revealed the presence of integrase genes in all repeat copies, and DNA primases were also found in several of the unique concatemers as well (Fig. 5). A similar arrangement of genes, predicted repeats, and gene contents are known to occur in small mobile elements called phage-inducible chromosomal islands (PICIs). PICIs are sometimes found in genomic islands of cultured bacteria and can parasitize other temperate phages by interfering with and exploiting the temperate phage reproduction and DNA packaging machinery. Concatemers of PICI DNA reproduce post-excision by rolling circle replication and are believed to be packaged “by the headful” into the hijacked, infectious temperate phage capsids, replacing the native phage DNA (Penades and Christie 2015) with their own. By virtue of this phage hijacking, PICI mobile elements can increase their rates of horizontal transfer by as much as five orders of magnitude (Penades and Christie 2015; Martinez-Rubio et al. 2017; Fillol-Salom et al. 2018). The gene content, repeat size, and overall length support the hypothesis that the concatemeric reads discovered here by our nanopore sequencing method represent PICIs packaged into phage particles. In particular, the integrases and DNA primases we found in the concatemers are hallmark genes of PICIs. Further, the concatemeric whole number repeat copies and read lengths we observed were consistent with predicted PICI-like DNA synthesis and packaging strategies. Previously, these unique mobile phage parasites have been identified mainly in cultures of gram-positive and gram-negative bacteria (Chen and Novick 2009; Martinez-Rubio et al. 2017; Fillol-Salom et al. 2018). To the best of our knowledge, these results represent some of the first observations of concatemeric PICIs packaged in phage particles from natural samples.

**Figure 5.**
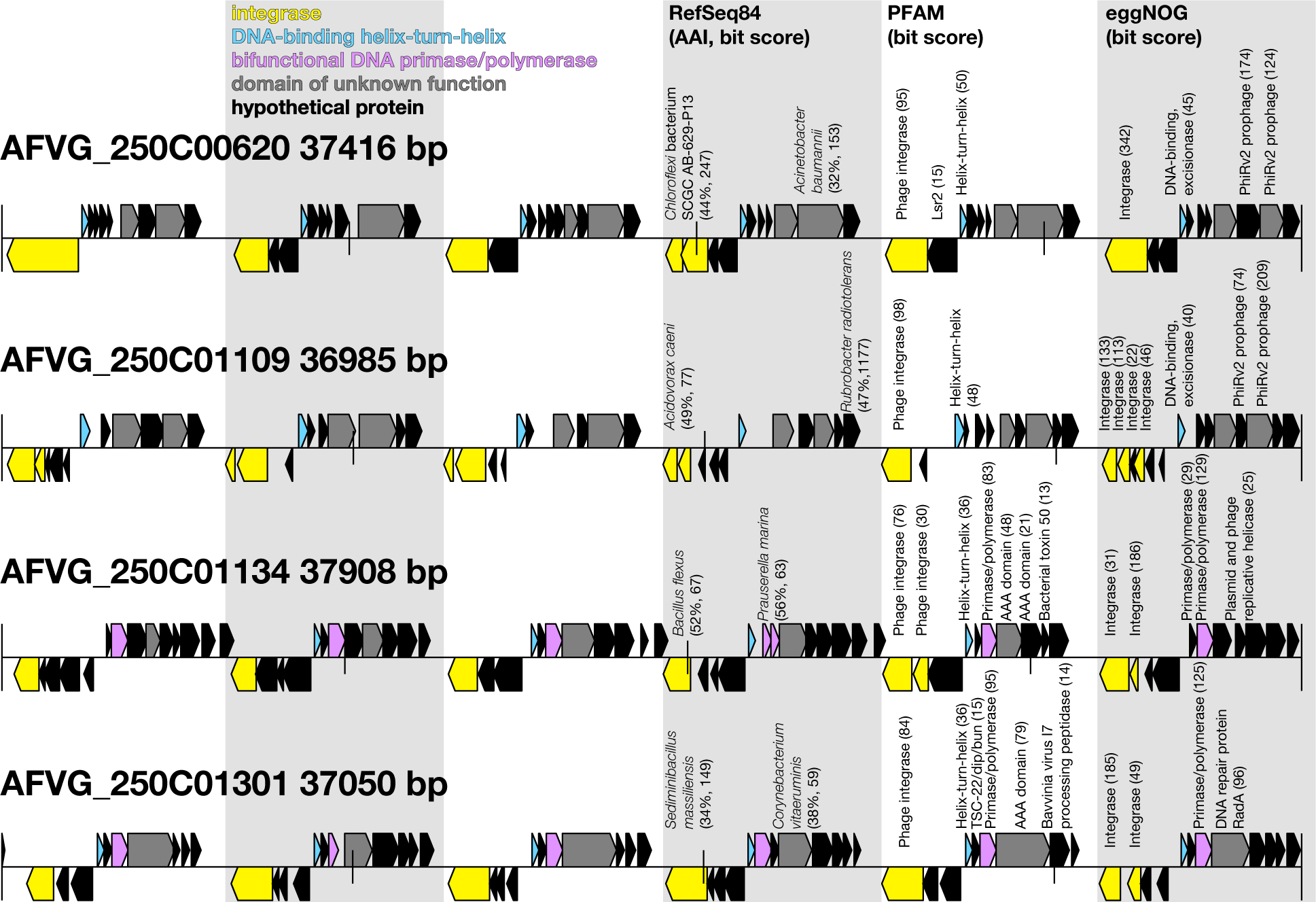
Structure putative phage-packaged PICI-like concatemer. Protein annotations from four representative concatemeric sequences recovered from the 250 m sample. Background shading represents repetitive regions. Protein coding sequences are color-coded based on functional annotations on PFAM (El-Gebali et al. 2019) (bit score >30). Taxonomic annotations are shown with top hit to organism of RefSeq84 (O’Leary et al. 2016), amino acid identity (AAI), and bit score. Functional annotations are shown with top hits and bit scores to protein domains in PFAM and eggnog (Huerta-Cepas et al. 2015).

## Discussion

Generation of dsDNA virus genome fragments from natural populations using standard, short-read metagenomic assembly techniques has enhanced understanding of wild virus populations, but the approach has inherent limitations. These limitations include difficulties in fully assembling or distinguishing highly repetitive sequence regions or repeats, an inability to accurately identify viral DTR sequences, and the inability to differentiate highly related genetically similar population variants. Our results show that many of these difficulties can be overcome by implementing a nanopore long-read sequencing strategy, which can yield thousands of high-quality, full-length virus genomes from a single sequencing run.

Further refinement of our approach has potential to improve with respect to the method’s efficiency, as well as the quantity, diversity and size range of virus genomes recovered. Examples include improving upstream virus purification, lowering the required DNA inputs, and expanding bioinformatic strategies for virus identification. Our initial approach required a starting input of roughly one microgram of virus-enriched high molecular weight DNA, but lowering the DNA input requirement should be achievable with future methodological refinement. Additionally, our current bioinformatic pipeline discarded many reads by stringently requiring that the primary k-mer binned sequences contain a tight size clustering, as well as the presence of DTRs to be considered as a virus genome. Viral sequences lacking a DTR on their termini, viral genome fragments of unequal lengths, or viral genotypes that simply were rare in the sample (so their bin sizes were lower than our cutoff threshold) would be omitted in our method as currently implemented. These challenges can be addressed in part by improving upstream virus purification methods, by using different and more inclusive strategies to identify viral sequences among all sequences, and by further incorporating long-read assembly strategies or polishing with short reads using other sequencing platforms, to better leverage the full range of virus genome fragments sequenced, beyond the fully intact, single read AFVGs described here.

A distinct advantage of our method over standard short read metagenomic approaches was the efficient recovery of full-length genome sequences without requiring any assembly. The size of genome sequences obtainable by our method is limited primarily by the quality and intactness of input DNA, along with current practical limitations of nanopore-based sequencing approaches (Tyler et al. 2018). Our approach also recovered full-length sequences spanning highly repetitive regions, that would otherwise collapse into single sequences as a consequence of the short-read sequence assembly process. The DTR regions flanking virus genomes, as well as the PICI-like virus-genome sized concatemer sequences we discovered specifically demonstrate these advantages. Our nanopore metagenomic sequencing method also has potential to capture highly variable genomic island regions in natural samples that would be difficult to assemble due to their inherent variability in sympatric populations (Thompson et al. 2005). Further applications should advance a deeper understanding of dsDNA virus genomic structure and variability, their fine scale population biology, and the prevalence of viral parasites like PICIs in the complex natural microbial assemblages.

## Methods

### Virus particle collection and DNA extraction

Two different approaches were tested for concentrating virus particles and enriching them from co-occurring microbial cell biomass in seawater samples. In the first approach, seawater was collected from 15 m depth using a Niskin bottle rosette and filtered through a 0.2 μm Supor PES Membrane Disc filters (Pall, USA). The resulting virus-enriched < 0.2 μm filtrate was collected onto a 25 mm, 0.03 μm Supor PES Membrane Disc filters (Pall, USA) to obtain virus particles. Filters were placed in 300 μl RNALater preservative (Ambion, Grand Island, NY) and stored at −80°C until final DNA extraction. In the second approach for concentrating and enriching for viral particles, 100 L of seawater was collected from 117 m depth and 250 m depths, then concentrated by tangential flow filtration (TFF). The seawater was first pre-filtered by peristatic pumping through a 0.1 μM Supor cartridge filter (Acropak 500, Pall, USA), and next concentrated by TFF over a 30 kDa filter (Biomax 30 kDa membrane, catalogue #: P3B030D01, Millipore, USA). Virus-containing retentates were stored at 4°C until final DNA extractions. DNA extractions for all samples were performed using Qiagen Genomic-tip 20/G (Qiagen, Hildern, Germany) following the manufacturers recommendations. See Supplementary Methods for extended details.

### ONT Sequencing

A total of 1 μg of purified DNA from each sample was used to prepare sequencing libraries using ONT’s standard ligation sequencing kit LSK109 (Oxford Nanopore, UK). The standard protocol was followed with the exception of the DNA repair and end-prep incubation step which was extended to 20 min. Care was given to minimize shearing when handling DNA. Sequencing was conducted on GridION X5 with FLO-MIN106 (R 9.4.1) flowcells (Oxford Nanopore, UK). Read basecalls were generated from the signal traces using Guppy v2.2.2.

### Preparation of short read libraries and Illumina sequencing

A total of 60 ng of genomic DNA from each sample was sheared to an average size of 350 bp using a Covaris M220 Focused-ultrasonicator (Covaris, Woburn, MA) with Micro AFA fiber tubes (Covaris, #520166, Woburn, MA). Libraries were sequenced using a 150 bp paired-end NextSeq High Output V2 reagent kit (Illumina, FC-404-2004, San Diego, CA). Illumina short reads were assembled into contigs in two steps. First, low quality sequence was removed using iu-filter-quality-minoche from the illumin-utils package (Eren et al. 2013). Second, remaining reads were assembled into contigs using the “meta-sensitive” mode of MEGAHIT (Li et al. 2015).

### Phage discovery bioinformatic pipeline

We developed a new assembly-free bioinformatic pipeline to isolate and polish full-length phage genomes from nanopore reads (Fig. 1). A coarse taxonomic annotation was first performed using Kaiju (Menzel et al. 2016) to bin the reads into two partitions (Table 1): one for reads with a cellular annotation (e.g. prokaryotic and eukaryotic) and another for the remaining reads (e.g. viral and unclassified). Subsequent read binning and genome polishing for each depth sample was conducted separately within read partitions.

To avoid any unnecessary bias, the next steps in the analyses were conducted independently of reference databases. In the initial binning step, 5-mer frequency vectors were calculated for all reads longer than 15 kb. Such 5-mer frequency features have been shown to be useful for binning and classifying prokaryotic genome sequencing data, using approximations of genome-wide patterns of k-mer usage (e.g. from assembled contigs or long reads) (Teeling et al. 2004; Saeed et al. 2012; Laczny et al. 2014; Beaulaurier et al. 2018). Due to the small genome size of marine bacteriophage (30-70 kb) it was possible to completely encompass an entire virus genome in a single read. Since such whole genome reads are not sampled from different regions of the genome, the only source of within-genome read variation in k-mer frequency vectors is due to the sequencing error, rendering the k-mer frequency vector a simplified representation of the genome sequence. Next, 5-mer frequency vectors were reduced from 512 to 2 dimensions using the uniform manifold approximation and projection (UMAP) technique (McInnes et al. 2018) and bins were automatically called in the resulting two-dimensional space using the HDBSCAN algorithm (McInnes et al. 2017) (Table 1).

Subsequently, read length distributions were constructed for each 5-mer frequency bin and used to identify bins that might contain full-length viral genome reads. Any bin with an anomalous enrichment of reads of a specific length, relative to the read length distribution of all reads in all bins, was marked as potentially containing full-length viral genome reads. Within each bin identified by this approach, reads were error-corrected using Canu (Koren et al. 2017), and any error-corrected reads within the anticipated viral genome length range (Steward et al. 2000; Holmfeldt et al. 2013) were retained (Table 1). These error-corrected, putative full-length viral genome reads were clustered within each bin at >95% pairwise average nucleotide identity (ANI) (Table 1). One read was selected from each bin cluster to serve as the draft genome reference, and the remaining reads in the cluster were used for polishing. Only polished reference sequences possessing the direct terminal repeat (DTR) structures typical of many dsDNA tailed bacteriophage (Casjens and Gilcrease 2009) were retained.

### K-mer binning of nanopore reads

Reads were annotated with Kaiju v1.6.2 (Menzel et al. 2016) using the options “-a greedy -e 5” and the NR+euk database (http://kaiju.binf.ku.dk/database/kaiju_index_nr_euk_2018-02-23.tgz). Reads with annotations for cellular organisms were partitioned, while the reads with viral annotations or lacking any annotation results were partitioned into a separate group. Although the initial purpose of this step was to serve as an *in silico* filter to further enrich for viral reads in the noncellular partition, subsequent analysis revealed the presence of significant amounts of phage DNA in the “cellular” read partition. We therefore separately submitted both read partitions to bioinformatic phage discovery pipeline.

Read binning was done by projecting reads into a 2-dimensional embedding of their 5-mer frequency vectors. Specifically, all 5-mers in read were counted and the counts of reverse-complement 5-mers were combined, resulting in a vector, ***Z***, for each read, *i*, denoted as ***Z***_***i***_ = (*Z*_*i,j*_, …, *Z*_*i,V*_), were *V* = 512 combined 5-mers. Following addition of a small pseudocount to ensure non-zero counts, the counts were normalized by the total number of 5-mer counts in the read and log2-transformed:

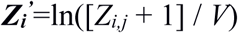

The resulting *N* × *V* matrix of log2-normalized 5-mer frequency, where *N* is the number of reads, was subjected to dimensionality reduction with UMAP v0.3.2 (McInnes et al. 2018) using the options *n_neighbors* = 15 and *min_dist* = 0.1. Bins were automatically called in the resulting 2-dimensional embedding with HDBSCAN v0.8.18 (McInnes et al. 2017) using the option min_cluster_size = 30 (Fig. 2a). All reads not assigned to a bin were omitted from further analysis.

### Isolation of bins containing reads spanning full phage genomes

Bins enriched for reads spanning an entire phage genome contained a significant enrichment of reads of a specific length, accounting for expected length variation resulting from indel sequencing errors. The probability density function (PDF) of the read lengths of all reads included in the analysis was subtracted from the PDF of read lengths in each bin. This curve representing the difference of PDFs was assessed using area under the curve (AUC) calculations along a sliding window of 1500 bp. Any bin for which the AUC in a 1500 bp region contained >= 4% of the total AUC across the full range of read length values between 20-80 kb was considered to have a set of read lengths significantly deviating from the background read length distribution (Fig. 2b). Such a deviation from the background read length distribution indicated the possible presence of many binned reads spanning full-length linear phage genomes. The expected size of the linear genome, *G*, could be estimated based on the location of the spike in the read length distribution.

### Clustering binned reads by average nucleotide identity

For each bin found to contain evidence of full-length phage genome reads, error correction was done with Canu v1.8 (Koren et al. 2017) using the options “maxThreads=4 maxMemory=12 genomeSize=*G* -correct corOutCoverage=400 stopOnLowCoverage=0 - nanopore-raw”, where *G* is estimated in the previous step. In each bin, the resulting corrected reads shorter than 90% of the estimated genome size were discarded from further analysis. The remaining error corrected reads were highly enriched for full-length genome reads. The pairwise average nucleotide identity (ANI) values of these reads were subsequently calculated with FastANI v1.1 (Jain et al. 2018) using the options “--fragLen=1500 --minFrag=10 -k 10”. Pairwise ANI values were hierarchically clustered with the cluster.hierachy.linkage function from the python package Scipy v1.1.0 using the Ward variance minimization algorithm. ANI clusters were then called using the function cluster.hierachy.fcluster such that each read in an ANI cluster has a cophenetic distance <= 1.0 (Fig. 2c). An ANI cluster was retained for further analysis if (1) the cluster contained >= 11 reads (one draft genome read and ten reads for polishing), (2) an average of pairwise ANI values in the cluster >=95%, and (3) an average read length > 28 kb. If these criteria were met, the read with the highest average intra-cluster ANI was designated to serve as the genome reference sequence and the remainder were designated for use in polishing the reference.

### Draft genome polishing and deduplication

Polishing was conducted using using both Racon (Vaser et al. 2017) and Medaka (https://github.com/nanoporetech/medaka). The first polishing step, which was iteratively done three times, used Minimap2 v2.15 (Li 2018) with “-ax map-ont” and Racon v1.3.1 with the options “--include-unpolished --quality-threshold=9”. The polished output of this step was further polished with Medaka v1.4.3 using the supplied model file *r941_flip_model.hdf5*. Any residual adapter sequences were then pruned from the polished draft genomes with Porechop v0.2.3 (https://github.com/rrwick/Porechop) using the option “--no_split”.

In order to ensure that each polished sequence represents a unique phage genome, the polished sequences were deduplicated twice. In the first instance, an all vs. all alignment of the polished sequences within each “cellular” and “noncellular” read partition of a given depth sample was done to eliminate redundant sequences. In the second instance, the polished phage sequences derived from the “noncellular” partition were aligned to those derived from the “cellular” partition. In each step, polished sequences were considered redundant if alignment by Nucmer v3.1 (Kurtz et al. 2004) using “--nosimplify” resulted in a hit with >=95% identity and covering >=95% of both polished sequences. In the case of redundant polished sequences, the sequence with the higher number of polishing reads was retained.

Finally, the coding sequences (CDS) of the polished and deduplicated draft genomes were annotated with Prodigal v2.6.3 (Hyatt et al. 2010) using the option “-p meta”.

### Identification of direct terminal repeats

Subsequences representing the first and last 20% of the draft genome sequence were aligned using Nucmer v3.1 (Kurtz et al. 2004) with default parameters. A draft genome is considered to have a direct terminal repeat (DTR) if an alignment is produced and (1) the aligned positions in the starting subsequence are within the first 200 bp, and (2) the aligned positions in the ending subsequence are with last 200 bp.

### Identification of linear concatemer reads

Subsequences of 3 kb were taken from the beginning of all reads >15 kb and aligned to their full-length reads using minimap2 v2.15 (Li 2018) with the option “-x ava-ont”. All reads where >90% of the 3 kb starting subsequence aligned at least twice on the full-length read were considered concatemeric. The concatemer repeat size was determined by taking the median of the differences between the consecutive starting alignment positions in the full-length read. Concatemer repeat copy numbers were calculated by dividing the full read length by the concatemer repeat size.

### Phage genome validation: taxonomic and functional annotations

Predicted proteins were taxonomically annotated using LAST v756 (Kielbasa et al. 2011) against the RefSeq84 database (O’Leary et al. 2016). Phage genomes were annotated at the genus level if they contain one, three, or five or more proteins with top hits to phages infecting the same bacterioplankton genus (Supplementary Fig. S8). Predicted proteins were functionally annotated using hmmsearch (Finn et al. 2011) against the PFAM-A v30 database (El-Gebali et al. 2019), and top hits at >30 bit score were retained (Supplementary Fig. S9). Predicted proteins from the four concatemeric sequences were also functionally annotated with the eggNOG database (Huerta-Cepas et al. 2015) and all functional top hits with any bit score were retained (Fig. 5).

### Characterization and comparative analyses of AFVGs

Known viral genes were placed in a single database from multiple sources: RefSeq release 84 (Brister et al. 2015), the Earth Virome (Paez-Espino et al. 2016) the Global Ocean Virome (Roux et al. 2016a), and three Mediterranean metagenomic viromes (Mizuno et al. 2013a; Mizuno et al. 2016; López-Pérez et al. 2017). Nucleotide gene sequences, predicted from AFVGs as described above, were compared to this database and also to all RefSeq genes using lastal v828 (Kielbasa et al. 2011). The highest scoring match was taken for each gene, and these matches were grouped by AFVG. For each AFVG the total fraction of genes with matches and the cumulative amino acid identity of matches were calculated (Fig. 3 and Supplementary Fig. S6).

### Phage genome relative abundances in short- vs. long-read sequences

Relative abundances of polished genomes were compared between short- and long-read sequencing, using normalized coverage for short reads, and relative number of reads in polished genome bin for long reads (Supplementary Fig. S4). BWA-MEM v0.7.15 (Li 2013) and msamtools (Arumugam et al.) was used to map and filter corresponding short reads to polished nanopore genomes at >95% average nucleotide identity (ANI) across >45 bp. SAMtools (Li 2009) was used to calculate nucleotides mapping to polished genome. Coverage was calculated for each genome by dividing nucleotides mapped to the length of the genome, and normalized coverage was calculated by dividing coverage to genome by summed coverage across all genomes in sample.

## Code availability

All custom scripts used to perform bioinformatics analyses are available from https://github.com/nanoporetech/marine-phage-paper-scripts.

## Data availability

All data sets presented in this paper have been deposited in the Sequence Read Archive under BioProject accession number is: PRJNA529454 and BioSample accession numbers: SAMN11262775, SAMN11267325, and SAMN11267326.

## Author Contributions

E.F.D. and M.P. conceived of the project. E.F.D. led the project. P.D. and E.F.D. collected the seawater samples, prepared the virus-enriched fraction, and extracted and purified the DNA. D.J.T., M.P., E.H., S.J. and X.D. coordinated and conducted ONT sequence generation and its preliminary quality checks and analyses. J.B. engineered, developed, and tested the bioinformatic pipelines. J.B., E.L. and J.M.E performed analysis of all sequence data sets. E.F.D., J.B., J.M.E. and E.L. wrote the manuscript. E.F.D., J.B., J.M.E., E.L., E.H., S.J. and D.J.T. contributed to the figures or to editing of the manuscript. This work was supported in part by a grant from the Gordon and Betty Moore Foundation (GBMF #3777 to E.F.D), and the Simons Foundation (#329108 to E.F.D).

