## Supplementary Methods, Figures, and Tables for "Assembly-free single-molecule nanopore sequencing recovers complete virus genomes from natural microbial communities": supplementary_methods_and_figs.pdf

### Supplementary Text

### Extended Methods

#### DNA Purification from 0.03 uM filters

Two different approaches were tested for concentrating virus particles and separating them from co-occurring microbial cell biomass in seawater samples. In the first approach, seawater from 15 m depth was collected in the North Pacific Subtropical Gyre (NPSG) during the Hawaii Ocean Experiment Legacy II cruise (KM1513; http://scope.soest.hawaii.edu/data/hoelegacy/). Seawater was collected from 15 m depth using a Niskin bottle rosette attached to a conductivity-temperature- depth (CTD) package (SBE 911Plus, SeaBird). For each sample, 2 L of seawater were filtered through a 25 mm, 0.2 µm Supor PES Membrane Disc filters (Pall, USA) housed in Swinnex 25 mm filter manifold via peristaltic pumping. The resulting virus-enriched < 0.2 um filtrate was subsequently filtered onto a 25 mm, 0.03 µm Supor PES Membrane Disc filter (Pall, USA) to obtain virus particles. Immediately after collection, all virus-containing 0.03 µm filters were placed in 300 ul RNALater preservative (Ambion, Grand Island, NY) and stored at -80°C until final DNA extraction. A total of 41 filters (representing 82 L of filtered seawater total) were extracted, and the resulting DNA pooled to yield the virus DNA for subsequent nanopore sequencing.

For DNA extractions, a modified Qiagen Genomic-tip 20/G (Qiagen, Hildern, Germany) was performed as follows: The 0.03 um filters containing phage particles from seawater were thawed on ice for 30 minutes, the samples were centrifuged 10,000 rpm for 60 seconds, and the RNALater (Ambion, Grand Island, NY) preservative was removed by pipette. Filters were then pooled into 5 mL sterile falcon tubes, 6 filters per tube.

An RNase A solution (200 ug/mL) was prepared by adding 20 ul of 10 mg/ml RNase A to 1 ml of Buffer B1 (50mM Tris•HCl, pH 8; 50mM EDTA, pH 8; 0.5% Tween-20; 0.5% Triton X-100). Next, 1 mL of the Buffer B1 lysis buffer containing RNase A was added to each tube of 6 pooled filters and inverted several times to coat the filters thoroughly. Then, 20 uL of a 100 mg/ml stock solution of lysozyme prepared in sterile water, and 45 uL of a Proteinase K solution (20 mg/mL) prepared in sterile water was added to each tube of 6 pooled filters, followed by gentle mixing and incubation at 37°C for 30 minutes. Next, 350 uL of Buffer B2 lysis buffer (3M guanidine HCl; 20% Tween 20) was added to each tube of 6 pooled filters, and the solution was mixed by inverting each tube several times, all tubes were then incubated at 50°C for 30 minutes.

The lysates (prepared from 6 filters) were pooled and loaded onto single Qiagen Genomic-tip 20/G column, and purification performed following manufacturers recommendations (Qiagen, Hildern, Germany). Each Genomic-tip 20/G column was equilibrated with 1 mL of Buffer QBT equilibration buffer (750 mM NaCl; 50 mM MOPS pH 7.0; 15% isopropanol; 0.15% Triton X-100) by gravity flow. Sample lysates were then mixed by inverting several times, and carefully pipetted sequentially onto one equilibrated Genomic-tip 20/G column, allowing samples to enter the resin by gravity flow. Next, 1 mL Buffer B1 was combined with 350 uL Buffer B2, and all sample lysate tubes were rinsed carefully with this 1 mL solution, and the rinse solution applied to the same Genomic-tip 20/G column. The Genomic-tip 20/G column was washed by gravity flow by applying 1 mL of Buffer QC wash buffer (1.0 M NaCl; 50 mM MOPS, pH 7; 15% isopropanol) three times, in succession. Finally, the genomic DNA was eluted from the column by two successive applications of 1mL Buffer QF elution buffer (1.25 M NaCl; 50 mM Tris•HCl, pH 8.5; 15% isopropanol), resulting in a purified DNA preparation in a 2 mL final volume.

The column purified DNA was concentrated by isopropanol precipitation as follows: The DNA eluant was split into two 2 mL conical screw cap tubes, 1mL per tube. The DNA was precipitated by adding 0.7 mL of room-temperature isopropanol per each 1 mL of DNA solution, followed by mixing by gentle inversion. After 2 hours at room temperature, the DNA was pelleted by centrifugation at 10,000 x g for 30 min at 4°C. The supernatant was removed, and the DNA precipitate washed by gentle addition of 1 mL of cold 70% ethanol and incubation for 60 seconds, followed by centrifugation at 10,000 x g for 15 min at 4°C. The supernatant was removed, and the DNA pellet air dried for 10 min. The purified DNA next resuspended in a final volume of 12 uL of 1X TE buffer (10 mM Tris•HCl, pH 8.0; I mM EDTA pH 8.0), and allowed to dissolve for a minimum of 10 min at room temperature, before final storage at 4°C. DNA quality was assessed by spectrophotometry (Nanodrop) and agarose gel electrophoreses, and final yields assessed by Quant-iT Picogreen dsDNA fluorimetric assay (Invitrogen, ThermoFisher, USA; catalogue #P7589,).

**DNA Purification from virus-enriched tangential flow filtration retentates**

In a second approach for collecting viral particles, seawater samples were collected from 117 m depth (22° 1841 N, 157° 05.57 W on April 2, 2018) and 250 m depth (24.33.5438 N, 160.50.6582 W on April 8, 2018) on the same oceanographic expedition (http://scope.soest.hawaii.edu/data/falkor2018/falkor2018.html) and concentrated by tangential flow filtration (TFF). For each sample, 90 L of seawater was collected using a Niskin bottle rosette attached to a conductivity-temperature- depth (CTD) package. The seawater was pre-filtered by peristatic pumping through a 0.1 uM Supor cartridge filter (Acropak 500, Pall, USA). The resulting virus-containing filtrate was concentrated by TFF over a 30 kDa filter (Biomax 30 kDa membrane, catalogue #: P3B030D01, Millipore). Subsequently, the retentate from 100 L of < 0.1 um seawater was reduced to a volume of 200 mL, and the tangential flow filter was backflushed with 100 mL of permeate to release virus particles trapped in the filter and the concentrated viruses were recovered in a final retentate volume of 300 mL. The virus-containing retentates were stored at 4°C until final DNA extraction.

Immediately prior to DNA extraction, the 300 mL retentates were concentrated using a Millipore Centricon Plus-70 Centrifugal Filter Units (10 KDa) (catalog: UFC701008, MilliporeSigma) following the manufacturers recommended protocol, resulting in a volume of 2-3 mL. A second round of centrifugal concentration of the resulting concentrated retentate was performed on a 100K Microsep Advanced centrifugal concentration device fitted with an Omega Advance membrane (modified polyethersulfone; catalogue #MCP 100C41, Pall, USA) following the manufacturers recommended protocol, resulting in a final volume of 200 uL of concentrated retentate, which was used for final DNA purifications. Lysis and DNA purification were performed in a single tube using the Qiagen Genomic-tip 20/G protocol per manufacturer's recommendations as described above. DNA quality was assessed by spectrophotometry and agarose gel electrophoreses, and final yields assessed by Quant-iT Picogreen dsDNA fluorimetric assay (catalogue #P7589, Invitrogen).

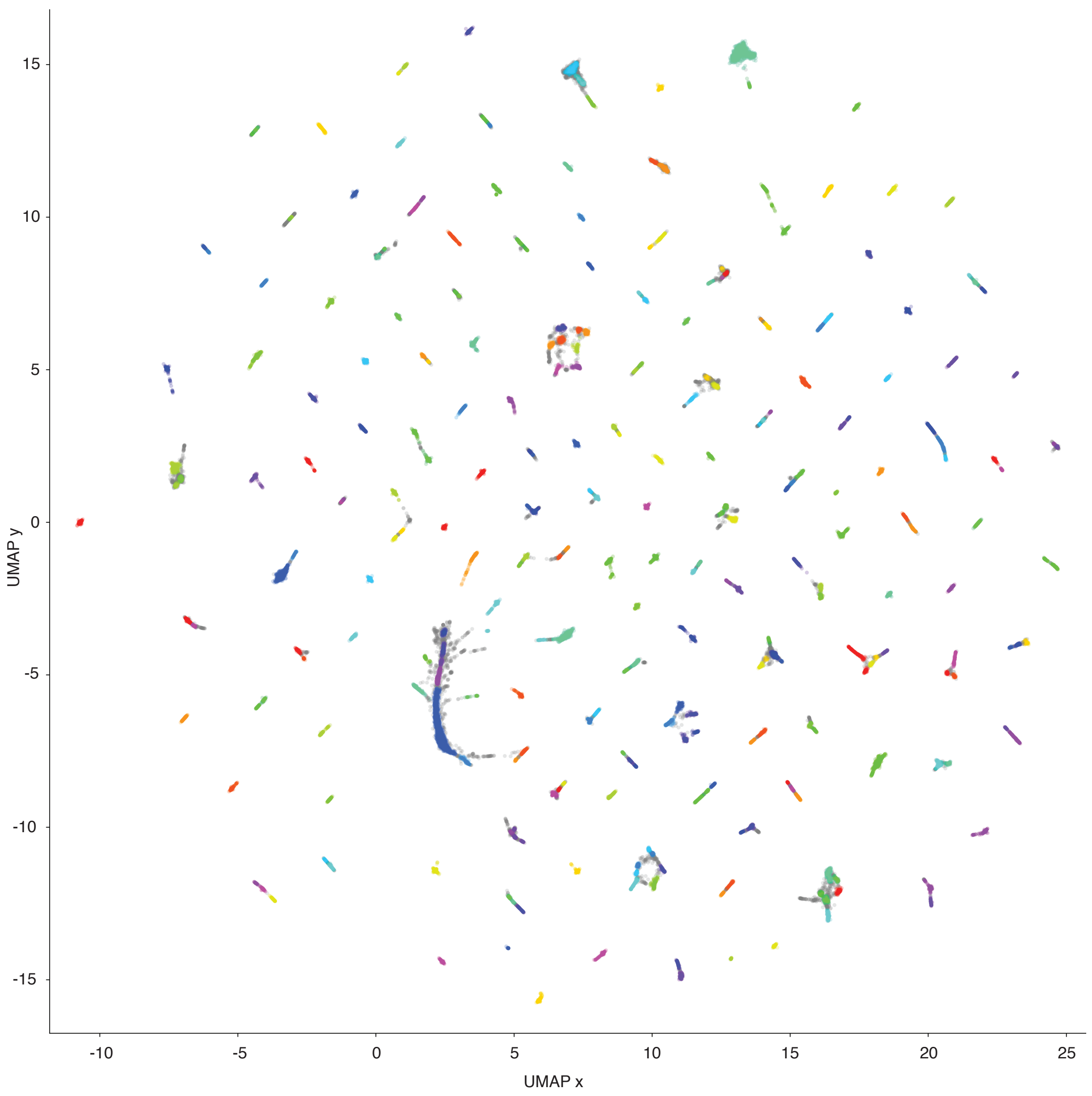


**Supplementary Figure S1. Nanopore reads simulated from the uvMED phage genome**

Nanopore reads simulated from the uvMED phage genome collection (Mizuno et al. 2016) (see Methods) were first size filtered to retain only reads > 15 kb, represented by their normalized 5-mer frequencies, and dimensionally reduced into a 2D embedding using UMAP (McInnes et al. 2018). Reads in the 2D embedding are colored based on their assignment to the 279 bins called by HDBSCAN (McInnes et al. 2017). Some bin colors are redundant due to the large number of bins. Reads not assigned to a bin are colored grey.

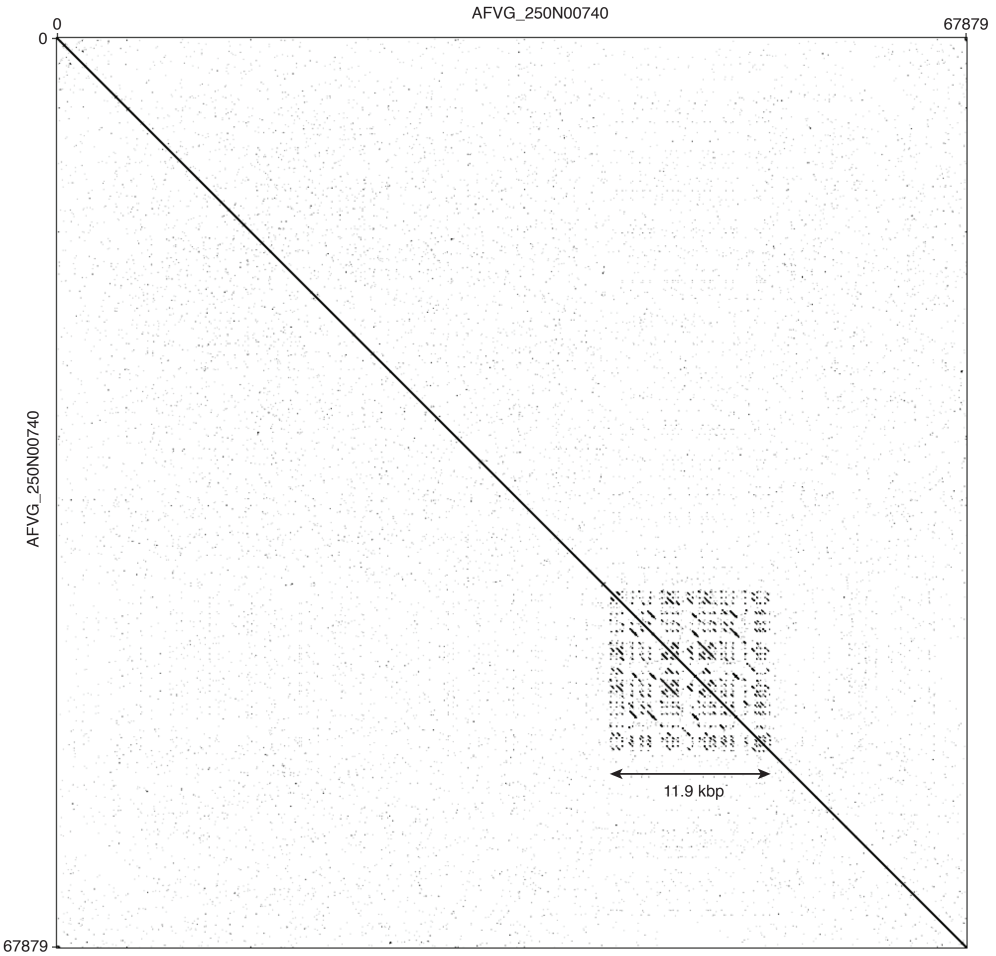


**Supplementary Figure S2. Dot plot showing complex repeats in a polished virus genome**

The 67.8 kb polished draft genome AFVG_250N00740 obtained from the 250 m sample reveals an 11.9 kb region of complex repeats that would likely present significant difficulties for short-read assembly tools.

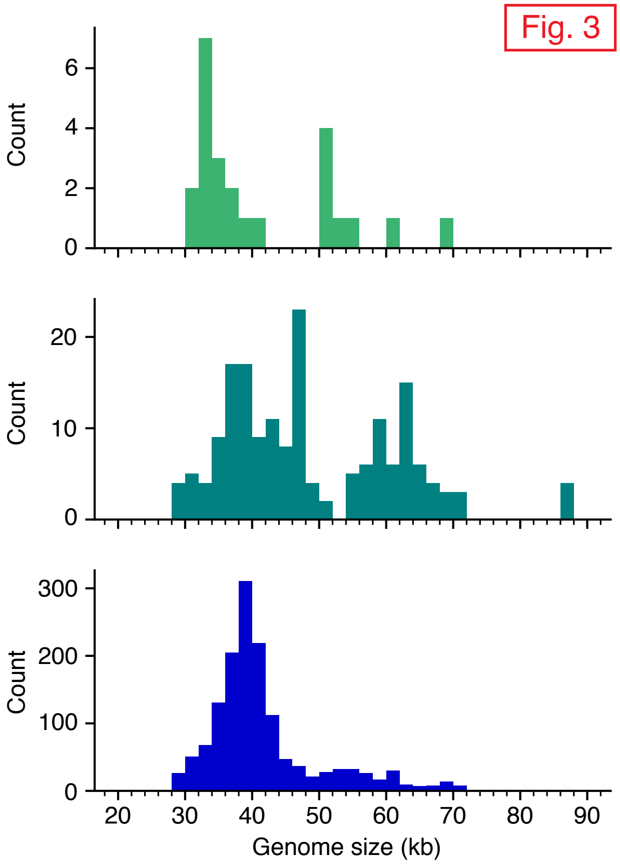


**Supplementary Figure S3. Polished draft genome yields and length distributions**

The number (counts) and genome lengths of the high-quality draft genomes obtained from the 15 m (top), 117 m (middle), and 250 m (bottom) seawater samples.

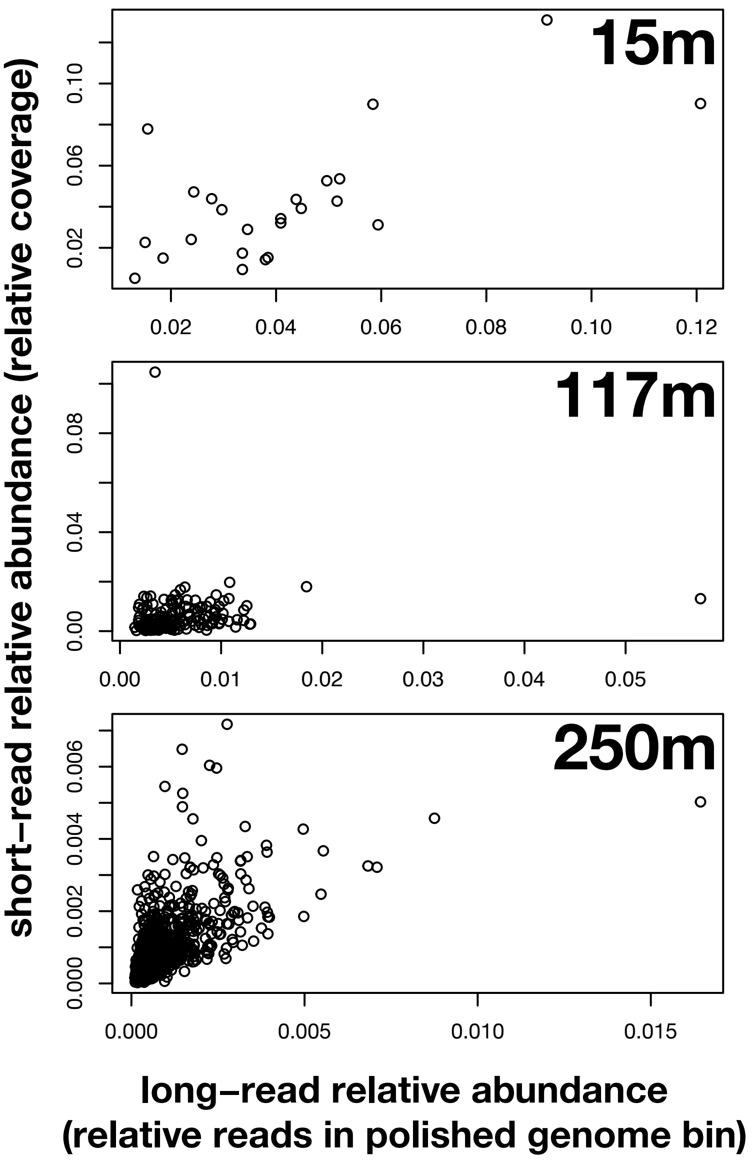


**Supplementary Figure S4. AFVG bin read abundances versus short read sequence coverage**

Each point represents one nanopore AFVG polished genome. X-axis represents normalized nanopore abundance, calculated by number of reads in an AFVG bin divided by total number of reads in all AFVG bins. Y-axis represents normalized Illumina short read abundance, calculated by coverage to an AFVG polished genome divided by total coverage to all AFVG polished genomes. bin read numbers versus Illumina read coverage.

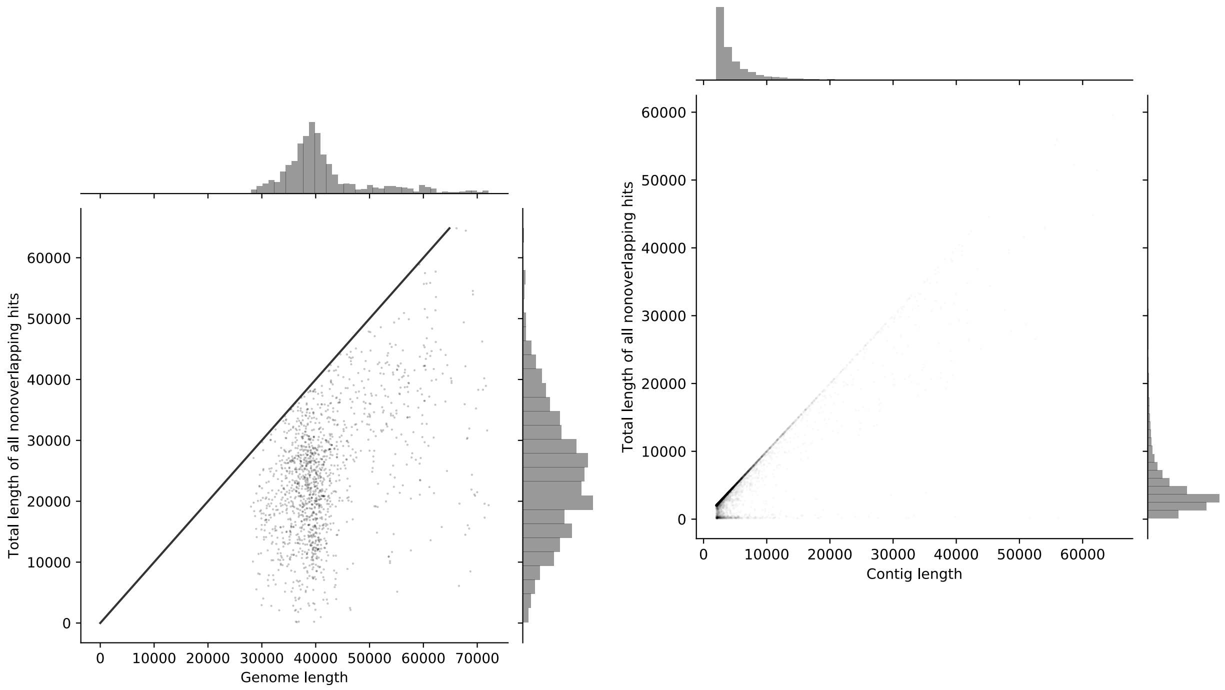


**Supplementary Figure S5. AFVG lengths versus short read contig coverages**

**Fig. S5a:** Each point represents a unique polished nanopore AFVG. The x-axis represents the AFVG genome length and the y-axis represents the total length of all non-overlapping matches from the short-read assembly contigs. A line is drawn showing the complete coverage boundary.

**Fig. S5b:** Each point on the plot represents a unique short-read assembly contig. The x-axis represents the short-read assembly contig length, and the y-axis represents the total length matched by nanopore AFVGs. (No line is drawn on the full coverage boundary because most points fall on that line).

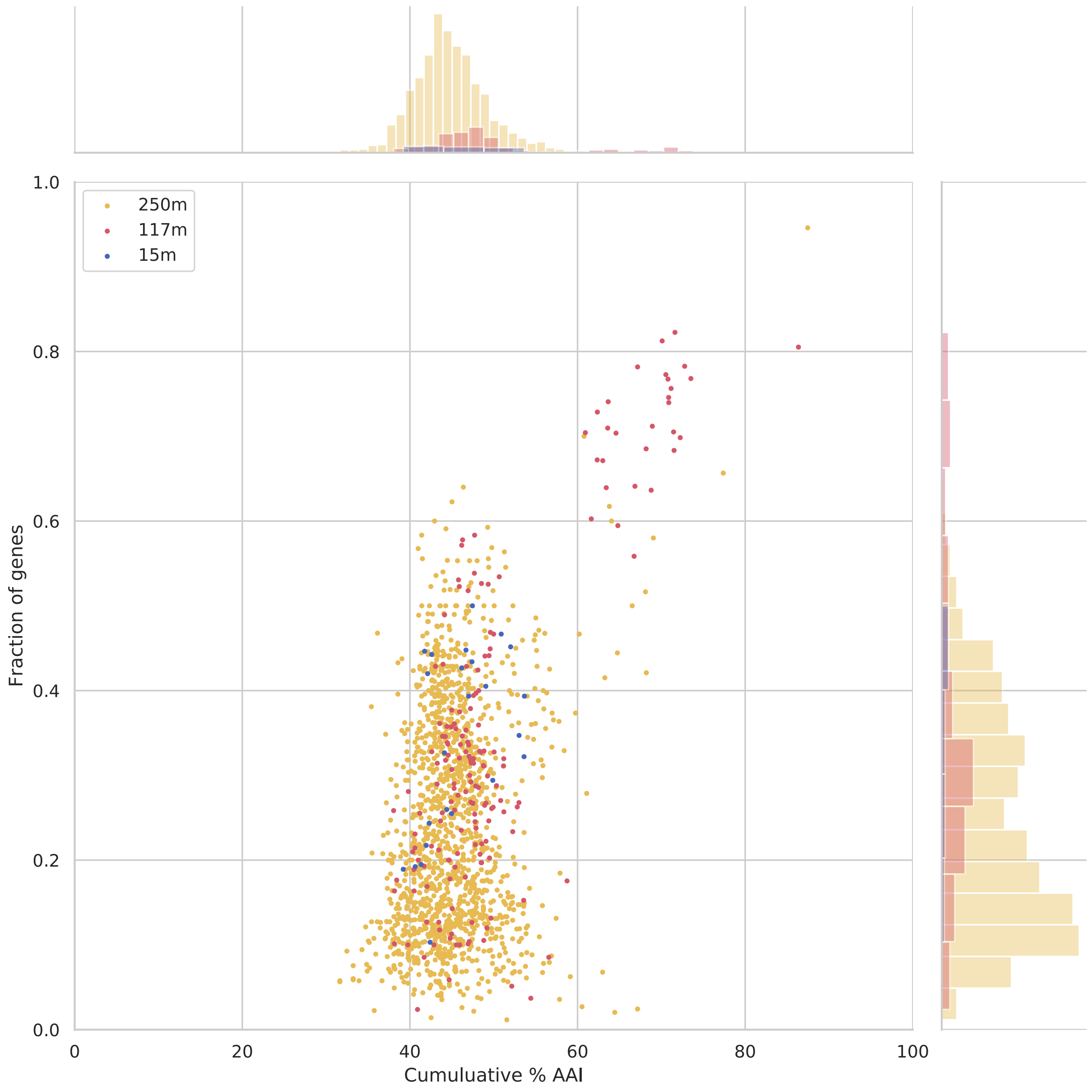


**Supplementary Figure S6. Similarity of AFVG genes to genes in the NCBI RefSeq database (release 84).**

Each point represents an AFVG colored by the depth at which it was found. The y-axis encodes the fraction of genes with matches to reference genes (y-axis), and the x-axis encodes the cumulative percent amino acid identity (%AAI) of matches to each AFVG. The marginal histograms show the distribution of values for "cumulative %AAI" (top) and "fraction of genes" (right) grouped by depth (Blue, 15 m sample; Red, 117 m sample; Orange, 250 m sample).

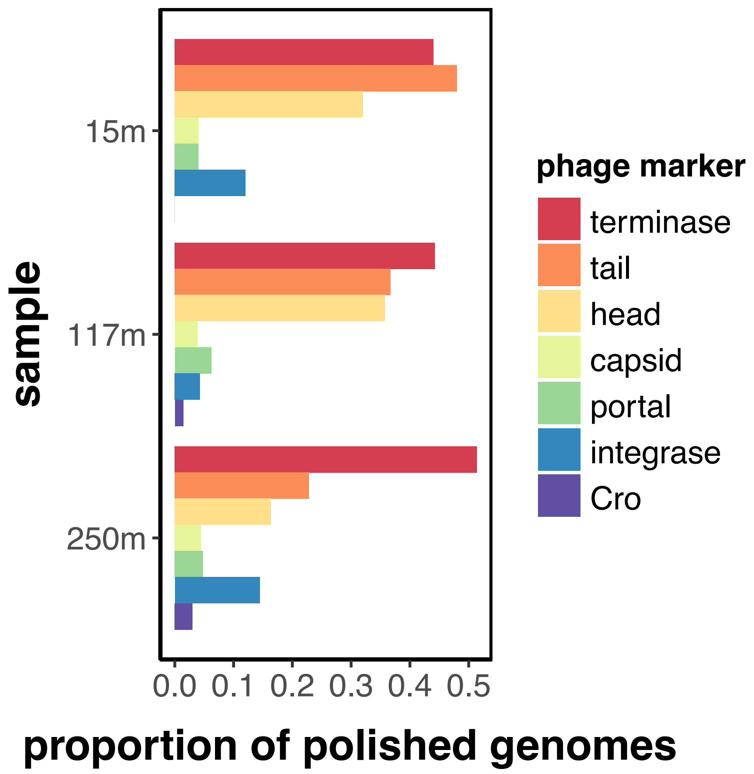


**Supplementary Figure S7. Virus marker genes found in AFVGs from each sample**

Proportion of polished nanopore AFVGs at each depth having one or more top protein hit (PFAM bit score >30) to the following virus and prophage marker proteins: terminase, tail, head, capsid, portal, integrase, Cro.

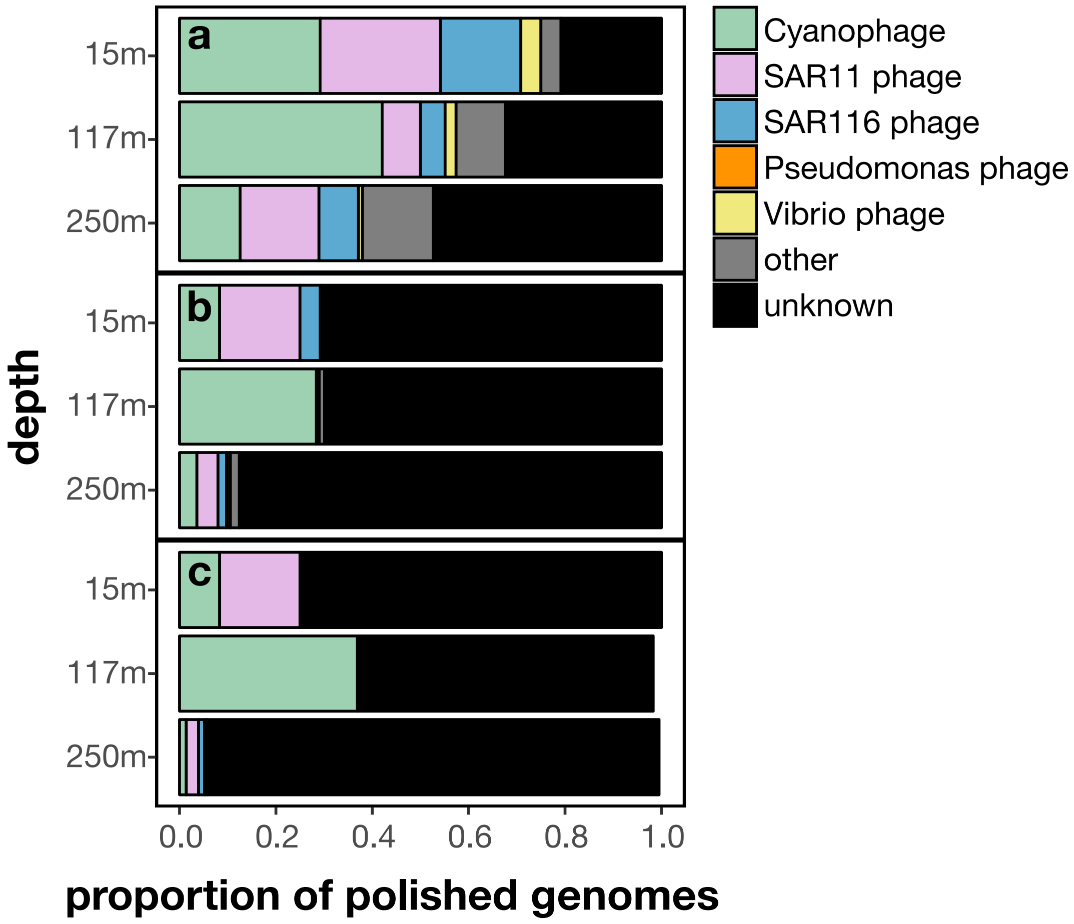


**Supplementary Figure S8. Putative virus taxon types identified in AFVGs in each sample**

Proportion of polished nanopore genomes functionally annotated to common virioplankton taxa on RefSeq84 (O'Leary et al. 2016) using cut-offs based on the following number of AFVG proteins in a single AFVG having a top match to a single genus: a. one or more proteins b. three or more proteins and c. five or more proteins.

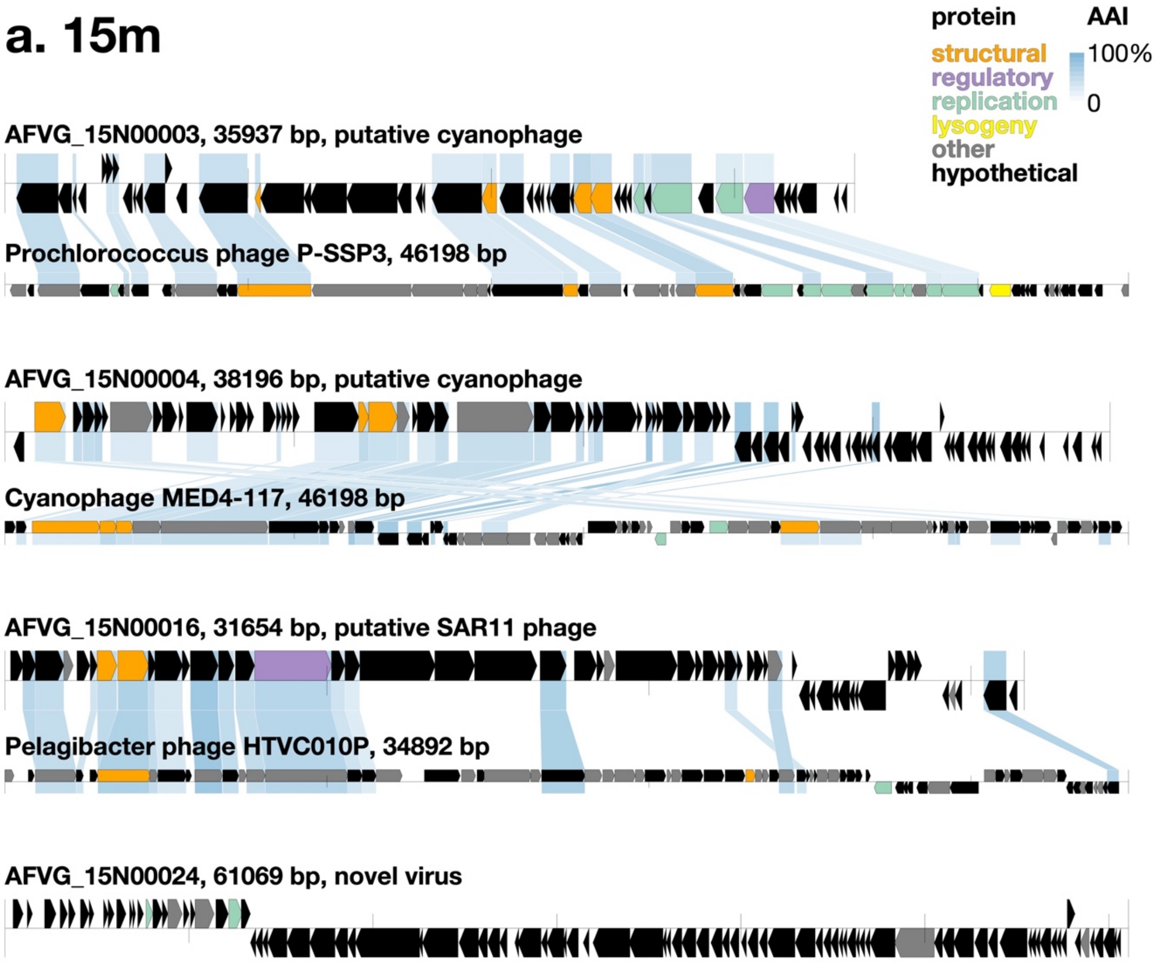


**Supplementary Figure S9a. Annotation plots of representative AFVGs.**

Figures show genomic structure and functional annotations of four abundant AFVGs recovered from each sample: a. 15m, b. 117m, and c. 250m. Protein coding sequences are color-coded based on functional annotations to PFAM (El-Gebali et al. 2019) (bit score >30). For AFVGs with three or more top hits to the same genus on RefSeq84 (O'Leary et al. 2016), the genome of their top hit is included below for reference, with blue shading representing amino acid identity (AAI) between homologous proteins.

## b. 117m

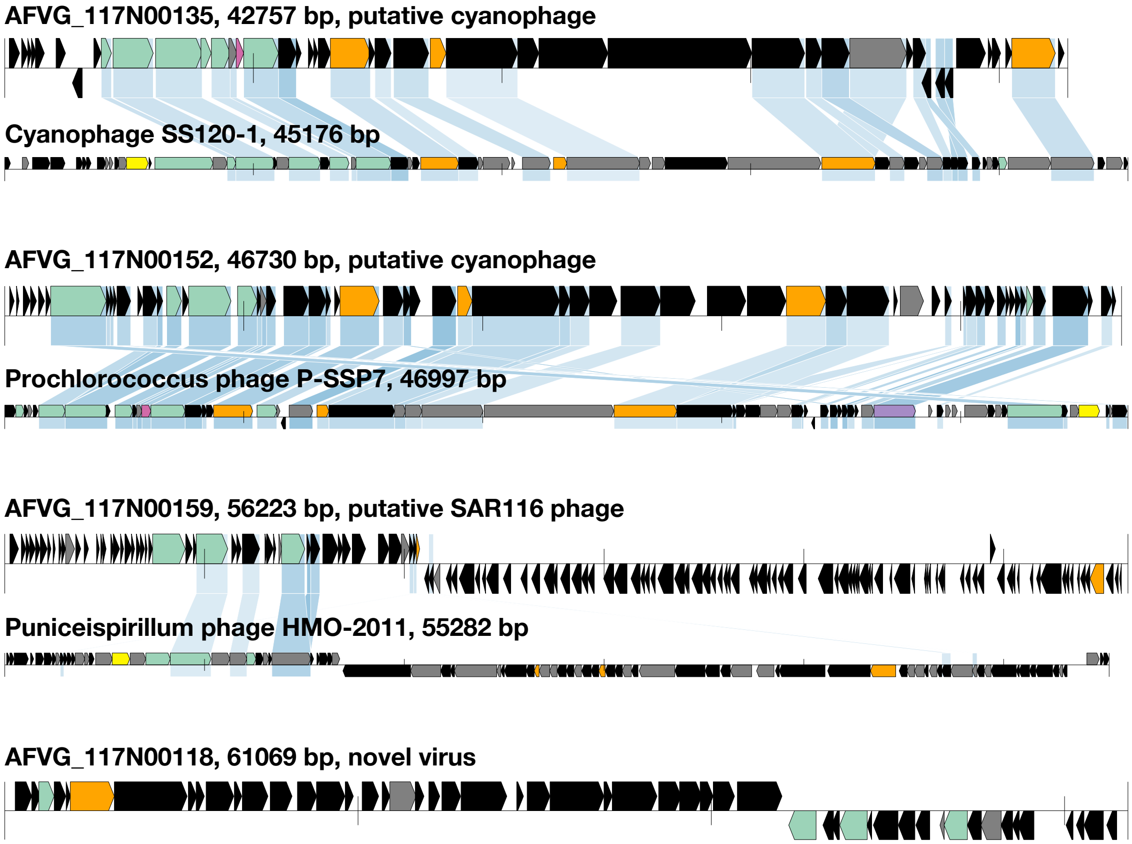


**Supplementary Figure S9b. Annotation plots of representative AFVGs.**

Figures show genomic structure and functional annotations of four abundant AFVGs recovered from each sample: a. 15m, b. 117m, and c. 250m. Protein coding sequences are color-coded based on functional annotations to PFAM (El-Gebali et al. 2019) (bit score >30). For AFVGs with three or more top hits to the same genus on RefSeq84 (O'Leary et al. 2016), the genome of their top hit is included below for reference, with blue shading representing amino acid identity (AAI) between homologous proteins.

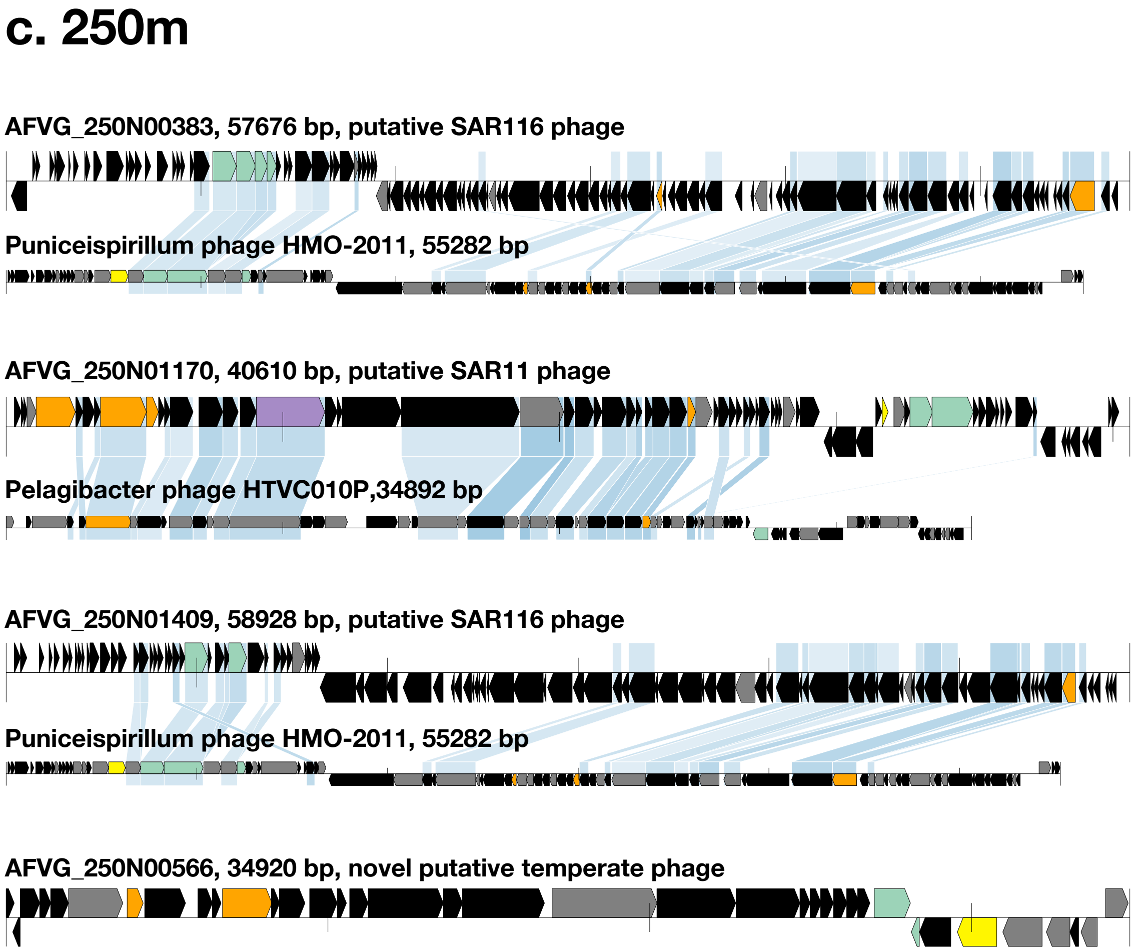


**Supplementary Figure S9c. Annotation plots of representative AFVGs.**

Figures show genomic structure and functional annotations of four abundant AFVGs recovered from each sample: a. 15m, b. 117m, and c. 250m. Protein coding sequences are color-coded based on functional annotations to PFAM (El-Gebali et al. 2019) (bit score >30). For AFVGs with three or more top hits to the same genus on RefSeq84 (O'Leary et al. 2016), the genome of their top hit is included below for reference, with blue shading representing amino acid identity (AAI) between homologous proteins.

**Supplementary Table 1. Statistics for 1630 polished genomes obtained from different samples**

Genome details, direct terminal repeat (DTR) size, GC content, and coding sequence (CDS) information for each polished draft genome are indicated. Each genome was polished from the specified sequencing read.

**Supplementary Table 2. Statistics for 11 polished linear concatemer sequences obtained from the 250 m sample**

Concatemer details, GC content, and coding sequence (CDS) information for each polished draft genome are indicated. Each genome was polished from the specified sequencing read.
